# Rethinking global carbon storage potential of trees. A comment on Bastin et al. 2019

**DOI:** 10.1101/730325

**Authors:** Shawn D. Taylor, Sergio Marconi

## Abstract

**Key Message:** Bastin et al. 2019 used flawed assumptions in calculating the carbon storage of restored forests worldwide, resulting in a gross overestimate.

**Context:** Bastin et al. 2019 use two flawed assumptions: 1) that the area suitable for restoration does not contain any carbon currently, and 2) that soil organic carbon (SOC) from increased canopy cover will accumulate quickly enough to mitigate anthropogenic carbon emissions.

**Aims:** We re-evaluated the potential carbon storage worldwide using empirical relationships of tree cover and carbon.

**Methods and Results:** We use global datasets of tree cover, soil organic carbon, and above ground biomass to estimate the empirical relationships of tree cover and carbon stock storage. A more realistic range of global carbon storage potential is between 71.7 and 75.7 GtC globally, with a large uncertainty associated with SOC. This is less than half of the original 205 GtC estimate.

**Conclusion:** The potential global carbon storage of restored forests is much less than that estimated by Bastin et al. 2019. While we agree on the value of assessing global reforestation potential, we suggest caution in considering it the most effective strategy to mitigate anthropogenic emissions.

## Main

Bastin et al. (2019) (hereafter referred to as Bastin 2019) use a novel machine learning based method to model global tree canopy cover potential. After accounting for current tree canopy cover and areas already occupied by urban and agricultural land they estimate 900 Mha of potential tree canopy cover available worldwide for reforestation. Using biome specific estimates of Tonnes C/ha they calculate the global carbon storage potential of this 900 Mha of tree canopy cover. The Tonnes/C ha-1 values for each biome are derived from average estimates of total carbon storage from two studies of forest (Pan et al. 2011) and tropical grassland (Grace et al. 2006) carbon stock. Thus from their calculation a hectare of restored tree canopy is equivalent to adding a full hectare of carbon stock potential regardless of the vegetation already in place, and results in an overestimate of the global carbon stock potential of restored trees.

To better estimate the relationship between total carbon stock density and tree cover we randomly sampled locations from four global datasets of 1) above ground biomass (Woods Hole Research Center 2019), 2) soil organic carbon (SOC) to 1-meter (Hengl et al. 2017), 3) percent tree cover (Hansen et al. 2013), and 4) the corresponding biome (Olson et al. 2001). We further subset these locations to those within protected areas (Levels I-V, UNEP-WCMC and IUCN (2019)) to minimize human influence on vegetation development and better represent the full carbon storage potential. Across all biomes there is already ample carbon stock at all levels of tree cover, and the relationship is weak in several biomes due to the contribution of SOC (Fig. 1). The slope of this relationship is a more accurate representation of the potential carbon stock gained with tree cover. For example in Tropical Grasslands Bastin 2019 estimate that an additional 0.5 ha of canopy cover (an additional 50% canopy cover) will add 141.25 Tonnes C. The empirical relationship shows an additional 50% tree cover in this biome means an additional 25.6 Tonnes C/Ha on average. Further, the Boreal Forest and Tundra biomes have a negative relationship between carbon stock and tree canopy cover, potentially resulting in a net carbon source if tree canopy cover was added in these biomes. Applying the updated estimates across all 14 biomes results in 28.4 GtC of potential carbon stock if the additional 900 Mha of global tree canopy potential was realized, and 71.7 GtC if the negative contribution from Boreal and Tundra biomes are removed.

**Figure 1:**
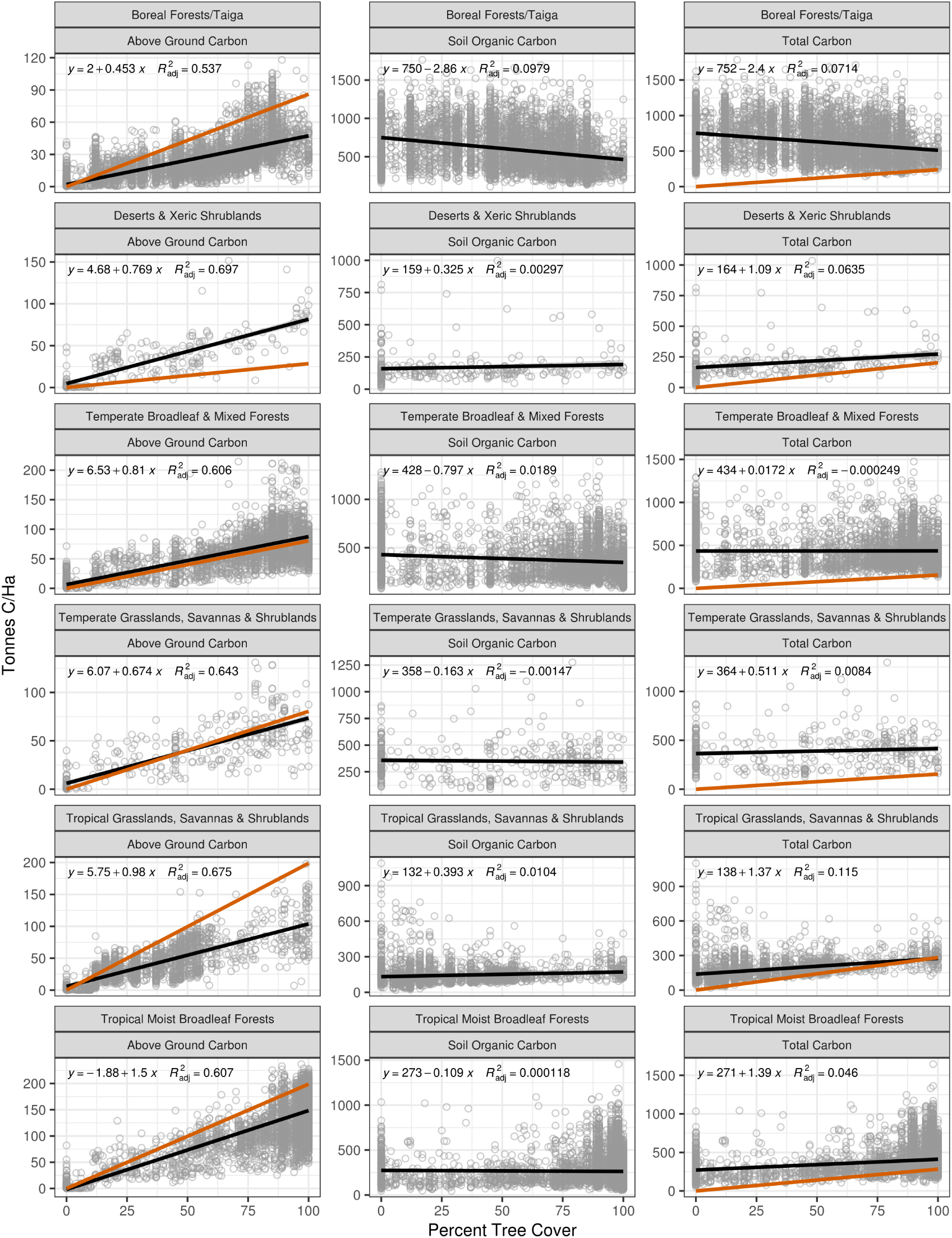
The relationship between carbon stock and tree cover for 6 of the 14 global biomes using global datasets (black regression line and grey points). The red lines for Total Carbon indicate the assumed increase in Tonnes of C/Ha for every increase in tree cover in the original analysis, while the red lines in Above Ground Carbon represents the original estimates minus the fraction of soil organic carbon. The global datasets were randomly sampled for land points within protected areas globally and querying the above ground biomass, 1-meter soil organic carbon, percent tree cover, and the corresponding biome. Above ground biomass was converted to carbon stock by multiplying by 0.5. Total carbon is above ground carbon plus soil organic carbon for each queried point. Note the difference in scales of the y-axis. See Figure S1 for relationships of all 14 biomes.

This calculation is further complicated by SOC. SOC makes up the majority of carbon stock in all biomes, and in seven biomes it has no relationship with tree cover (*p > 0.05*, see Supplemental Fig. S1). In boreal regions (the biome for 19.8% of the potential canopy area estimated by Bastin 2019) afforestation can cause a temporary increase of greenhouse gas emissions due to quicker SOC mineralization, which can take several decades to recover (Karhu et al. 2011). SOC also forms at rates of less than 0.5 Mg Ha^−1^ year^−1^ in many areas (Trumbore and Harden 1997, Gaudinski et al. 2000, Lichter et al. 2008), though sometimes up to 1.5 Mg Ha^−1^ year^−1^ (Shi and Han 2014), and it is unreasonable to assume increased tree cover would lead to SOC accumulation at a rate quick enough to effectively mitigate carbon emissions (He et al. 2016). To explore the potential carbon storage of increased global tree cover without considering the complexities of SOC we adjusted all estimates by removing the contribution of SOC. For the Bastin 2019 estimates we re-calculated the carbon stock potential minus the SOC fraction using the original sources (Grace et al. 2006, Pan et al. 2011). For our own estimates, we considered only aboveground carbon and its slope with respect to tree cover. With these estimates the global carbon storage potential is 104 GtC using the re-calculated estimates from Bastin 2019, and 75.7 GtC using the empirical relationships from the global datasets.

Bastin 2019 state that global tree restoration is “the most effective solution” for mitigating climate change. This conclusion uses simple assumptions which ignore complex carbon dynamics, potential feedback loops, societal costs, and carbon saturation as forests mature (see de Coninck et al. (2018) sec. 4.3.7.2 and references therein). For example, some authors consider afforestation and reforestation as an effective mitigation solution only in the tropics since it would reduce albedo in high latitudes (Fuss et al. 2018). Yet, increasing forested areas in the tropics would compete for agriculture and other land use, triggering a number of socio-economic impacts (Fuss et al. 2018). It is also difficult to place the 205 GtC estimate in the context of other mitigation options without a quantitative estimate of the timescale of global forest regrowth, which requires local studies using more nuanced analysis of carbon uptake (eg. Requena Suarez et al. 2019). Several other comments to Bastin 2019 have raised similar concerns. Namely that the original analysis does not adequately consider SOC, currently in place vegetation, or feedback loops such as fire and changed albedo (Friedlingstein et al. 2019, Lewis et al. 2019, Veldman et al. 2019). Veldman et al. (2019) re-analyzed the Bastin 2019 results using literature derived values of carbon storage and arrived at a potential 107 GtC from the original 900 Mha of canopy cover. Here, by using biome specific empirically derived relationships of carbon storage and canopy cover from global datasets, we show the potential global carbon storage of restored forests ranges between 71.7 - 75.7 GtC, less than 40% of the original estimate. Along with the other comments this demonstrates that the original Bastin 2019 estimate was clearly overestimated.

**Table 1:**
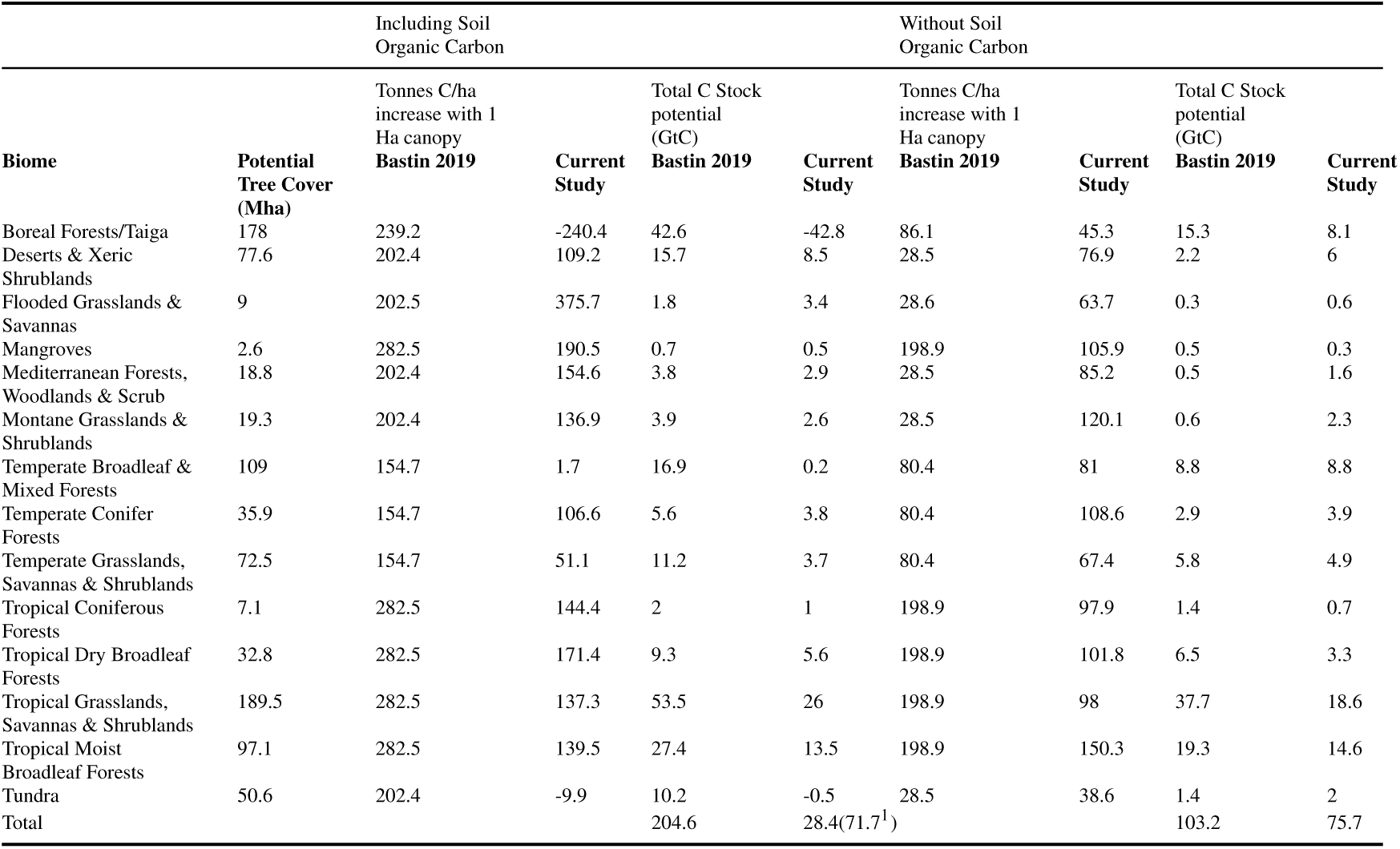
Estimates of the Tonnes C/ha relationship and per biome estimate of total carbon storage potential using the original estimates from Bastin 2019, estimates derived using global datasets in the current study, and all estimates adjusted to exclude soil organic carbon. The biome specific potential tree canopy cover is from Bastin 2019 Table S2. ^1^ 71.7 GtC is the global potential is calculated without considering Boreal Forests or Tundra, as these biomes have a negative relationship between total carbon stock and tree canopy cover.

## Supplemental Material

Supplemental Figure S1, carbon stock relationships for all 14 biomes. All code and extracted data is archived on Zenodo (https://doi.org/10.5281/zenodo.3364028).

## Acknowledgments

This research was supported by the Gordon and Betty Moore Foundation’s Data-Driven Discovery Initiative through Grant GBMF4563 to E.P. White.

