## Supplementary figures and images for "Rethinking global carbon storage potential of trees. A comment on Bastin et al. 2019"

### Supplemental Figure S1

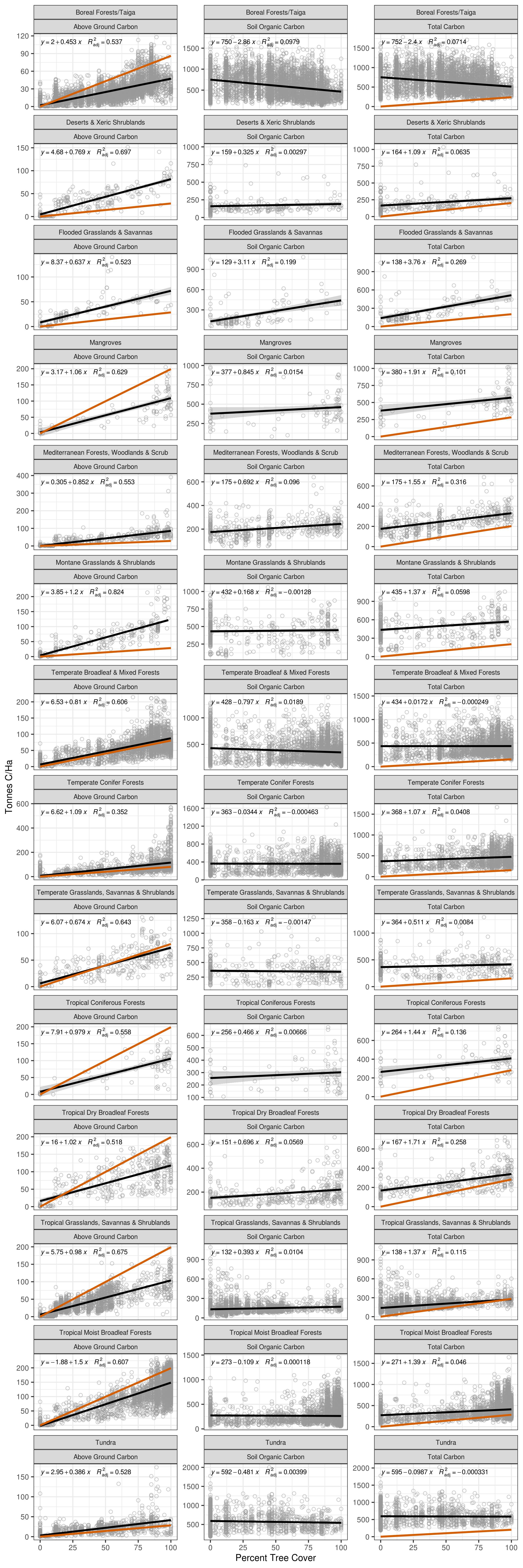
